# Closing the gaps: testing the efficacy of carbapenem and cephalosporins in treating late-stage anthrax

**DOI:** 10.1101/2024.09.21.614292

**Authors:** Assa Sittner, Elad Bar-David, Itai Glinert, Amir Ben-Shmuel, Josef Schlomovitz, Haim Levy, Shay Weiss

## Abstract

Anthrax is a fatal zoonotic disease caused by exposure to *Bacillus anthracis* spores. Treatment of systemic anthrax is usually efficient when using the right antibiotics as close as possible to exposure, preferably prior to symptoms’ onset as post exposure prophylaxis (PEP). The efficacy decreases as treatment is initiated later in disease progression. The CDC in its guidelines divides anthrax treatment to three different indications according to the progression of the disease: PEP, systemic and systemic with indications of CNS infection. While the prognosis of PEP or early treatment of systemic anthrax is very good, ingress of the bacteria into the CNS significantly decreases treatment efficacy, creating a substantial clinical challenge. Since anthrax in humans is rare, the CDC recommendations are mainly based on animal model experiments and data obtained from patients infected with other pathogens. Here we use rabbits to test the efficacy of the combined treatment of Meropenem and Doxycycline which is the first choice in the CDC recommendations for treating systemic patients with indication of CNS infection. In addition, we test the efficacy of the first-generation cephalosporin, cefazolin, in treating the different stages of the disease. We found that the combination of Doxycycline and Meropenem is highly effective in treating rabbits in our inhalation and CNS infection models. Cefazoline was efficient only as PEP or systemic stage treatment but not CNS infected animals. Our findings support the CDC recommendation of using a combination of Doxycycline and Meropenem for systemic patients with or without indications for CNS infection. We found that Cefazoline is a decent choice for PEP or early sage systemic disease but recommend considering using this antibiotic only if all other options are not available.

## Introduction

Anthrax is a zoonotic disease that effects mainly, but not solely, grazing animals [1, 2]. Natural transition to humans is rare and usually results form close contact with sick animals or contaminated animal products [3, 4]. Anthrax is caused by the spore forming, gram positive bacteria *Bacillus anthracis*. In most cases the spore is the infectious form. Inside the host, the spore germinates and in response to specific stimuli produces the two main virulence factors; the tripartite toxins and the capsule, encoded by the two virulence plasmids pXO1 and pXO2 [1, 2]. The toxins inactivate the host immune system, and the capsule protects the bacteria inside phagocytic cells and in addition has an immune-modulation effect. The tripartite toxin is composed of the lethal factor (LF), an endopeptidase specific for proteins of the MAP kinase pathway; edema factor (EF), a calmodulin dependent adenylate cyclase that disrupts cell cycle regulation and protective antigen (PA), that binds to a specific receptor, present on all cell types, forming a heptamer transporting the LF and EF into the cytoplasm. The *B. anthracis* capsule is a homo-polymer of γ-D-glutamic acid, that forms a negative charged helix that creating a barrier surrounding the bacterium [5, 6].

The type of disease in humans is determined by the route of infection; contact with skin lesions, injection, digestion or inhalation. Contact of spores or vegetative bacteria with compromised skin results in the most obvious and typical type of anthrax – cutaneous [1]. This form of the disease manifests in the painless formation of a typical lesion (eschar) surrounded by an inflamed region that in most cases recovers with minimal scarring even in the absence of antibiotic treatment. Fatality in this form of infection is between 5-30% and is the result of the transition of the infection from localized to systemic mode [7]. The resemblance of eschar to coal is the name source of the disease and bacterium (ἄνθραξ, anthrax in Greek) [3]. The second (and becoming the most common) form of infection is due to the digestion of contaminated meat from a near death animal, domestic or wild, resulting in gastro-intestinal disease [8]. This form of the disease is usually 100% lethal in the absence of antibiotic treatment and starts as a severe gastric inflammation and rapidly becomes systemic. Another form of this disease is the oropharyngeal were the infection causes the swelling of lymph-nodes in the throat that causes suffocation [9]. Since the accepted paradigm is that sporulation occurs postmortem as the contaminated blood is exposed to air [4], it is assumed that the vegetative form of the bacterium is responsible for this form of the disease. Deep tissue anthrax *as* a result injecting spore contaminated heroine uses was reported in the early 2010’s in Europe [10]. This massive tissue inflammation progressed, in the absence of antibiotic treatment, into systemic infection and death [11]. Unlike the cutaneous and oropharyngeal diseases, there are no new reports of the deep tissue form of the disease in the last decade. Though not considered as a natural disease [12-14], inhalation anthrax is the most worrying form of disease when *B. anthracis* spores use as a military or bio-terror weapon is considered [15-17]. Inhalation of spores results in a systemic disease without any early typical symptoms other than those resembling the a common flu [1]. The acute stage of the disease is short and is lethal even under antibiotic treatment [18]. Inhalation anthrax was considered an occupational disease that affected workers in goat hair processing mills [13]. Two major deadly events demonstrated the potential of using *B. anthracis* spores as a weapon; the Sverdlovsk accident [19-21] and the 2001 anthrax letter attacks [22, 23]. In Sverdlovsk, accidental spore discharge caused the death of people and farm animals up to 50 km from the source. In 2001, spores that leaked from sealed envelopes caused pulmonary anthrax in 11 people, several of them without a traced contact with the identified spore-containing articles [18, 23]. The systemic dissemination of the bacteria from the initial infection into the blood stream is common to all types of the disease. From the bloodstream the bacteria invade the internal organs and eventually crosses the blood brain barrier (BBB) causing meningitis and hemorrhage, most likely fatal [1, 2, 20, 24-27].

Treatment of anthrax patients is mainly based on antibiotics [28]. Currently the CDC treatment recommendations divides the treatment to Post Exposure Prophylaxis (PEP) and treatment of systemic (symptomatic) disease, depending on the stage of the disease [28]. Treatment of exposed non-symptomatic individuals (PEP} relays on oral administration of single antibiotic, usually ciprofloxacin of doxycycline. Treatment of symptomatic patients is more complicated and treatment with more than one antibiotic is recommended due to the possibility of meningitis, which is common at the late stages of the disease in humans and animal models. Since the data from actual patients is limited due to poor documentation and the low number of cases in the western world, most of the data concerning treatment efficacy is based on experiments in animal models. Previously, we and others demonstrated the efficacy of antibiotic treatment in different stages of the disease, from PEP through symptomatic animals to CNC infected animals [29-35]. In its 2023 guidelines, the CDC’s first choice for the treatment of symptomatic adults is the combination of meropenem and doxycycline (minocycline) [28]. While we tested meropenem as a single treatment in rabbits [30, 36], the combined treatment was not tested. In addition, though cephalosporins are not recommended for anthrax treatment, the CDC acknowledged the lack of experimental information regarding the efficacy of cefazolin, a first-generation cephalosporin, in treating anthrax [28]. Previously we demonstrated in Guinea pigs that PEP with Cefazolin resulted in full protection in the first 3 days, followed by the subsequent death of 50% of the infected animals, while under antibiotic treatment [29]. We proposed that though sensitive in vitro, *B. anthracis* expresses β-lactamases that enable the bacteria to grow under this antibiotic treatment. Herein we test the efficacy of cefazolin treatment of rabbits in different stages of the disease, PEP, different stages of the systemic disease and CNS infections. In addition, we test the efficacy of the combined treatment of high dose meropenem and doxycycline in treating rabbits in different stages of the systemic disease following airway infection.

## Results

### The efficacy of cefazolin as Post Exposure Prophylaxis

To test the effectiveness of cefazolin as PEP, rabbits were infected with virulent *B. anthracis* Vollum spores via intranasal instillation (IN) and treated 6h post-infection intravenous (IV) with 60 mg/kg cefazolin. The animals were then treated twice a day subcutaneously (SC) with the same dose for the total of 7 days. The animals were monitored for 14 days from the end of antibiotic treatment for signs of illness or death. 21 days from infection the animals were challenged SC with 100 LD_50_ of Vollum spores to test for any protective immune response that developed during the antibiotic treatment. The results of the experiment are presented in **Figure 1**. The result shows that all the animals (n=4) survived for 12 days while one rabbit succumbed to the infection 6 days from the end of the antibiotic treatment. The other 3 rabbits survived for 21 days. However, none of them developed protective immunity and they all succumbed to the SC challenge.

**Figure 1.**
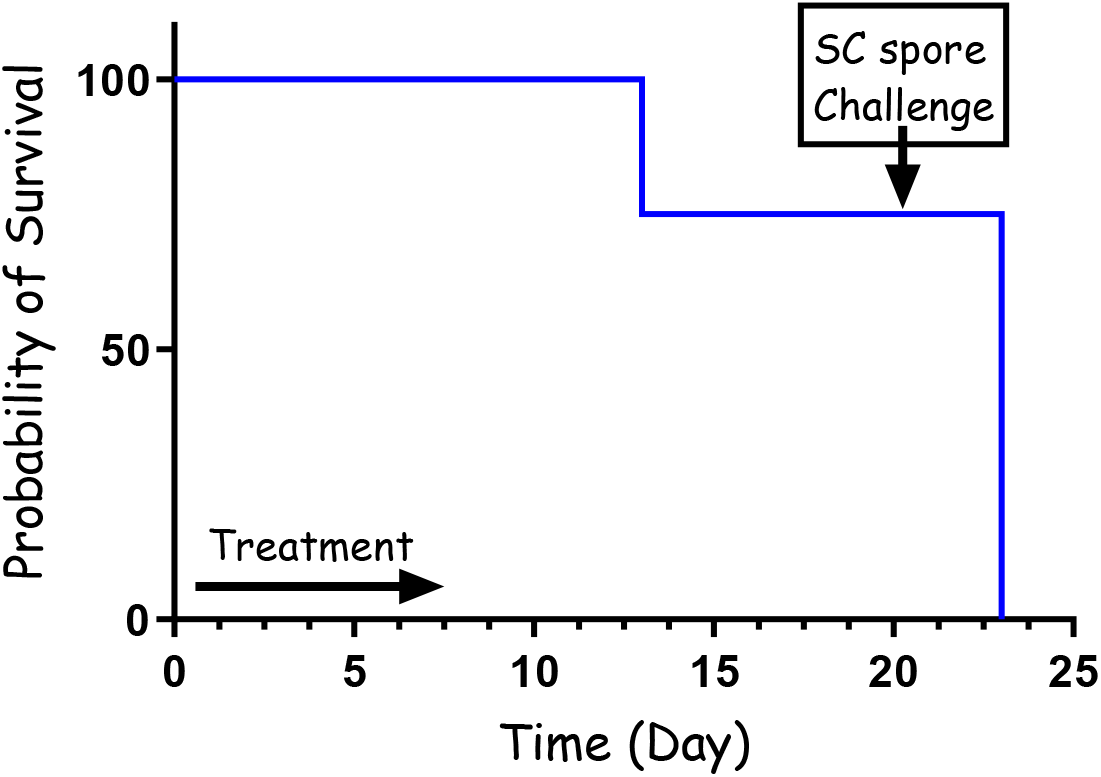
PEP treatment (60 mg/kg) with cefazolin following IN spore instillation of *B. anthracis* Vollum strain. 4 rabbits were sedated and infected IN with spores. Treatment was initiated 6h post infection, SC with 60 mg/kg cefazolin, twice a day. The treatment duration is marked by a vertical arrow and the SC spore challenge is marked with a horizontal arrow. Survival is presented as a Mayer Kaplan curve.

To further test the effectiveness of the cefazolin treatment, we repeated the PEP experiment with three major modifications. For this experiment the antibiotic dose was reduced from 60 mg/kg to 30 mg/kg, the first treatment was administrated 24h from the IN-spore instillation and the treatment lasted for a total of 5 days. The results presented in **Figure 2** demonstrate that the short, low dose treatment was as efficient as the full dose experiment. In this experiment, one rabbit succumbed to the infection while 7 rabbits were protected for 14 days from the IN infection. Furthermore, all the animals that survived post the antibiotic treatment developed protective immunity that protected them from a lethal spore challenge of 100 LD_50_, having survived for 14 additional days, after which they were euthanized.

**Figure 2.**
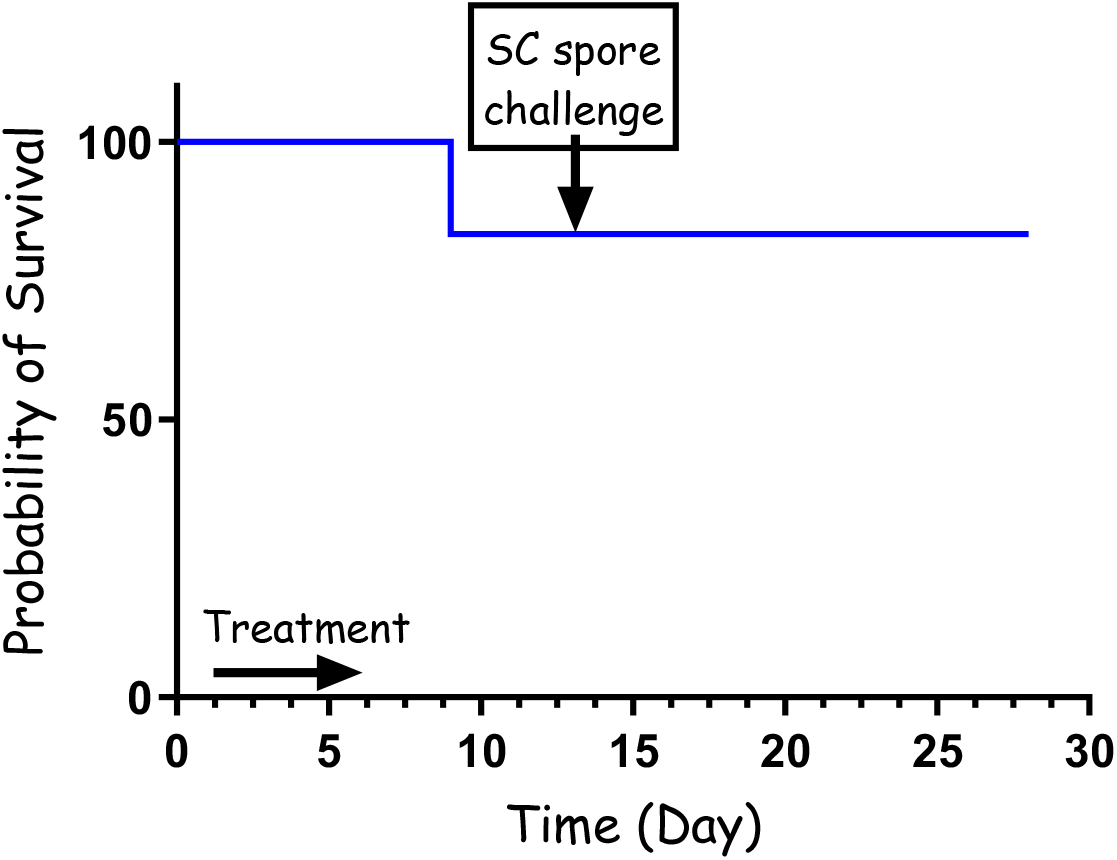
PEP treatment (30 mg/kg) with cefazolin following IN spore instillation of *B. anthracis* Vollum strain. 8 rabbits were sedated and infected IN with spores. Treatment was initiated 24 h post infection, SC with 30 mg/kg cefazolin, twice a day. The treatment duration is marked by a vertical arrow and the SC spore challenge is marked with a horizontal arrow. Survival is presented as a Mayer Kaplan curve.

### The efficacy of cefazolin as a treatment for systemic disease

To test the effectiveness of cefazolin in treating systemic disease, we infected the rabbits IN using our PennCentury device to spray Vollum spores intra-nasally as described previously. 20 h post infection, a blood sample was collected from each of the rabbits to determine the bacteremia level (CFU/ml), after which they were treated IV with 60 mg/kg cefazolin. The bacteremia level was determined by serial dilutions and plating, with the actual counting was performed the next day. At treatment initiation, 2 rabbits were below the limit of detection (10 CFU/ml), 5 rabbits were with bacteremia of 1×10^3^ to 2×10^4^ CFU/ml and one rabbit with bacteremia of 1×10^7^ CFU/ml (**Figure 3**). The rabbits were treated SC, twice a day for the total of 10 days with 60 mg/ml cefazolin and monitored for 21 days post infection. Cefazolin was effective in treating all the rabbits with bacteremia of up to 2×10^4^ CFU/ml and failed in treating the rabbit with 1×10^7^ CFU/ml (**Figure 3**). These results indicate that cefazolin is effective in treating rabbits with anthrax as PEP and in a systemic disease of up to at least 2×10^4^ CFU/ml.

**Figure 3.**
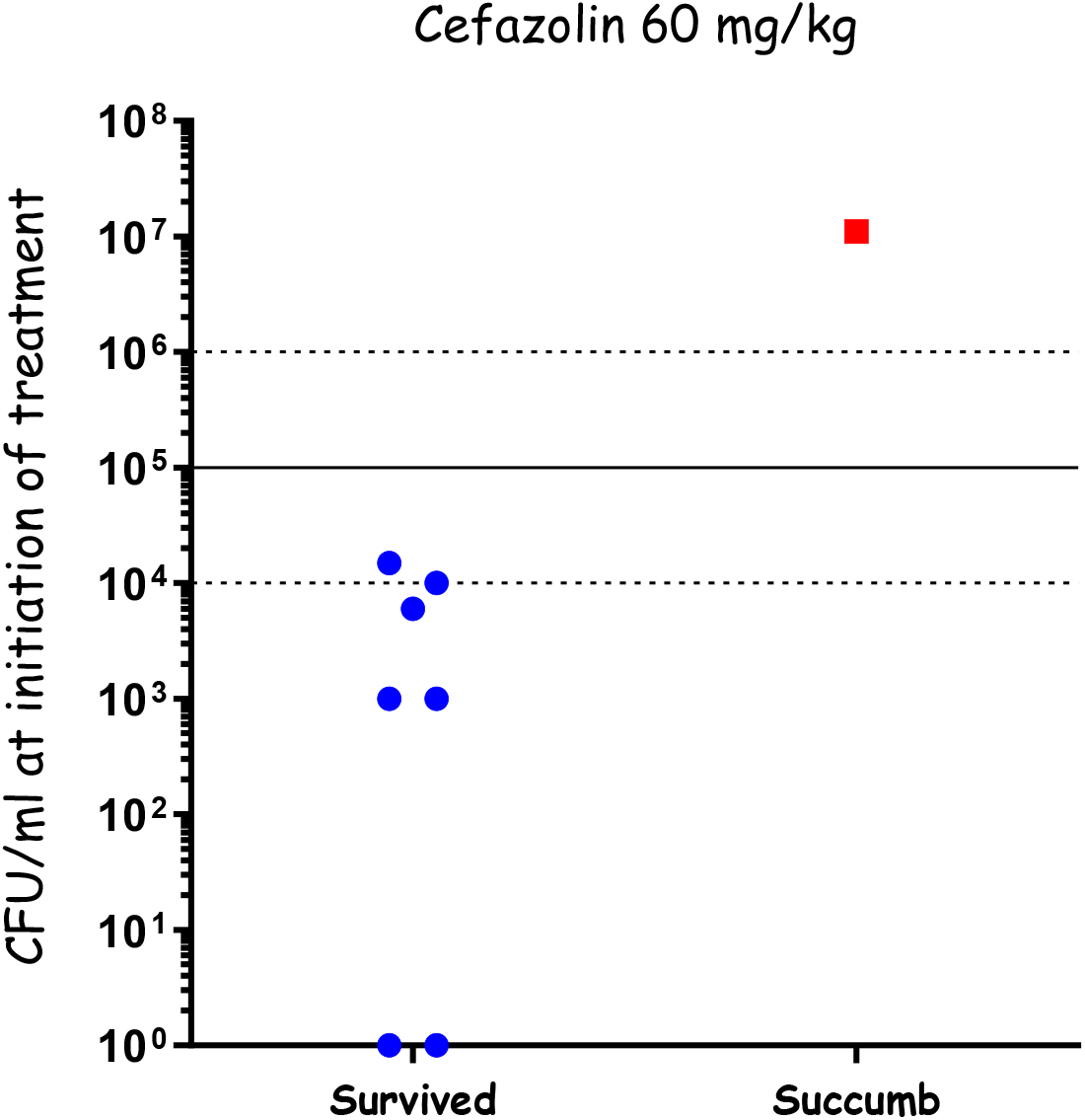
Cefazolin treatment of systemic disease following IN spray of *B. anthracis* Vollum spores. Rabbits were inoculated by spraying spores into the nasal cavity using a MicroSprayer ® Aerosolizer. Blood bacterial concentrations were determined at the time of treatment initiation. Survival (blue) or death (red) is presented in correlation to the initial bacteremia.

### The efficacy of cefazolin as a treatment for the meningitis stage of the disease

Treating anthrax meningitis is challenging. Previously we demonstrated that antibiotic treatment could protect rabbits from lethal CNS infections [30, 33, 36]. To determine the efficacy of cefazolin in treating anthrax related meningitis, we used our rabbit CNS infections model [36]. Rabbits were injected with 3×10^4^ CFU of vegetative encapsulated bacteria of the fully virulent Vollum strain. 6h post infection, antibiotic treatment was initiated. To ensure maximal efficacy we tested a regimen of a first dose of 200 mg/kg antibiotic administered IV, or combination of 60 mg/kg IV with 300 μl of 20 mg/ml administered ICM, directly into the CSF. In both cases the first treatment was followed by SC 60 mg/kg doses twice a day. None of the treatments were effective (**Figure 4**) as all of the infected animals died within 24h of the infection, indicating that cefazolin is not effective in treating the meningitic stage of the disease.

**Figure 4.**
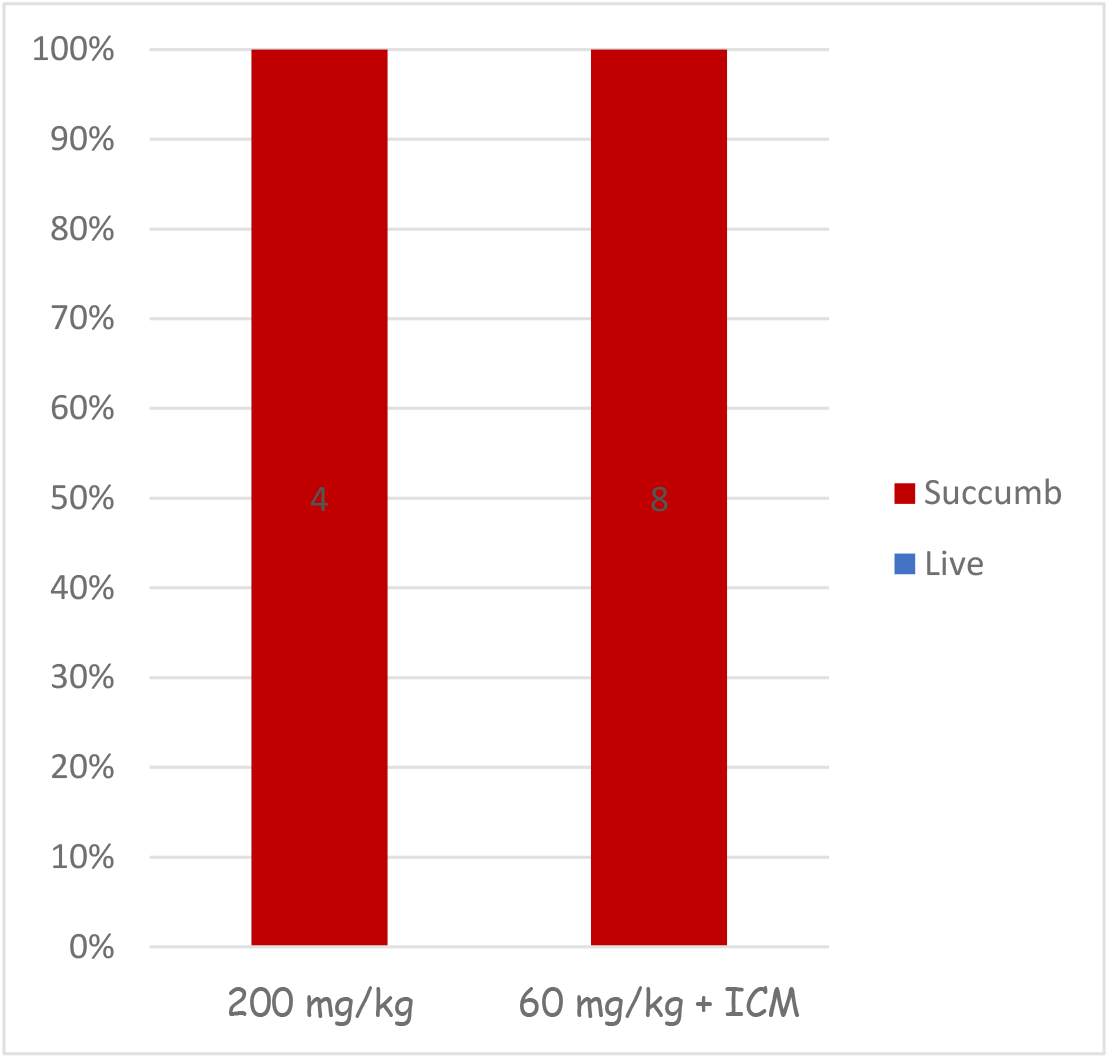
Cefazolin treatment of CNS infected rabbits. Groups of 4 or 8 rabbits were infected by injection of capsular vegetative Vollum bacteria into the cisterna magna and treated with cefazolin as indicated. Survival (blue) or succumbed (red) percent is presented.

### The efficacy of the CDC recommended treatment of systemic patient combining meropenem and doxycycline

The recent CDC guidelines for the treatment of adults with confirmed anthrax recommends a combined treatment of meropenem and doxycycline [28]. Previously we demonstrated that treatment with either meropenem (imipenem) or doxycycline as a mono therapy was effective in treating different stages of systemic anthrax [31, 33]. However, only high dose treatment of meropenem (150 mg/kg) was effective in treating anthrax meningitis in the rabbit CNS infection model [30, 36]. Therefore, we tested the efficacy of a combined administration of high dose meropenem (150 mg/kg) with doxycycline (15 mg/kg) to treat inhalation anthrax at different stages of the disease. Rabbits (n=17) were inoculated using the PennCentury device to spray Vollum spores as described previously [33]. 20 h post infection, blood a sample was collected from each of the rabbits to determine the bacteremia level (CFU/ml), followed by IV treatment of150 mg/kg meropenem and 15 mg/ml doxycycline. The variation in bacteremia was high and we treated animals with bacteremia levels in the wide range of <10 – 5×10^8^ CFU/ml, representing early to advanced stages of the disease (**Figure 5**). The combined treatment was highly effective, protecting all the animals with bacteremia of up to 1×10^6^ CFU/ml (n=10); 33% of the animals with bacteremia of 1×10^6^ to 1×10^7^ CFU/ml; and failed to protect the single animal with bacteremia of 5×10^8^ (**Figure 5**). Overall, this combined treatment was as affective as treating with doxycycline as a monotreatment indicating that there is no antagonism between these two drugs.

**Figure 5.**
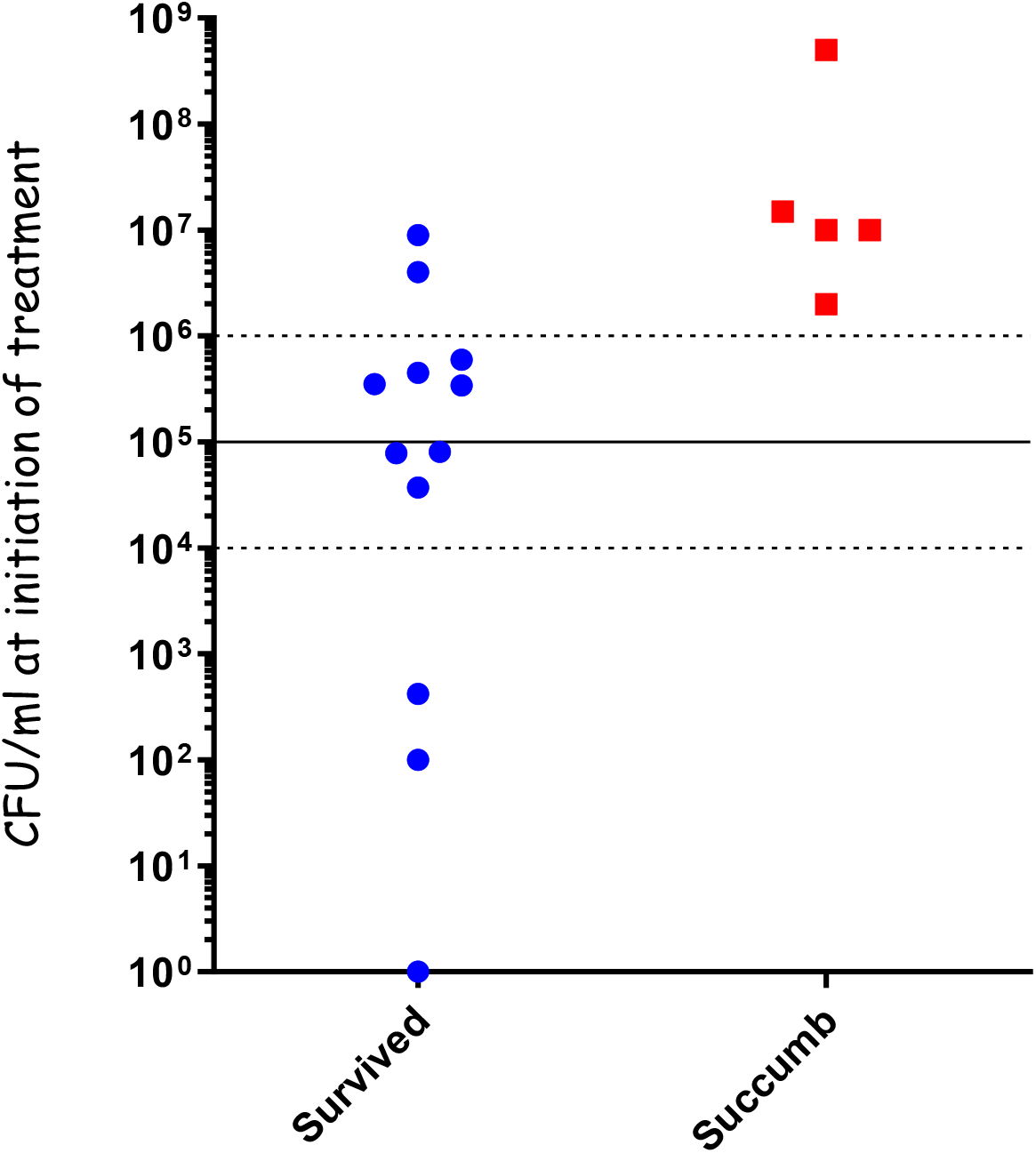
Combined treatment with meropenem and doxycycline of systemic disease following IN spray of *B. anthracis* Vollum spores. Rabbits were inoculated by spraying spores into the nasal cavity using a MicroSprayer® Aerosolizer. Blood bacterial concentrations were determined at the time of treatment initiation. Survival (blue) or death (red) is presented in correlation to the initial bacteremia.

## Discussion

Cephalosporins play a crucial role in the ICU setting, primarily due to their broad-spectrum antibacterial activity and relatively low toxicity [37]. as well as the fact that there are 5 generations of this drug that are specific for different types of bacteria [38]. During the anthrax letters attacks there were several cases in which patients were misdiagnosed and treated with cephalosporins and their condition deteriorated, resulting in several deaths [18]. Generation 1 is the most relevant to gram positive bacteria. *B. anthracis* is sensitive to this drug in vitro [29, 39]. In this manuscript, we tested in a rabbit model the efficacy of cefazolin, a generation 1 cephalosporin, in treating anthrax at different stages of the disease. As post exposure prophylaxis, we tested relatively short treatments of 5 to 7 days administrated twice a day with 60 mg/kg or 30 mg/kg. The efficacy of these treatments was relatively high, protecting in both experiments 85% of the animals (2/12) (**Figure 1 and 2**). The 60 mg/kg treatment, started 6h post infection was so effective that none of the surviving animals developed protective immunity, indicating that in these animals the spores did not have the opportunity to establish infection (**Figure 1**). On the other hand, the short treatment was not always affective, allowing relapse and development of lethal disease, as soon as 4 days after the end of the antibiotic treatment. A longer antibiotic treatment, such as the recommended 30 to 60 days, will probably prevent this relapse. Challenging this treatment by initiating the antibiotic treatment 24h post infection and using 50% of the dose (30mg/kg) for 5 days results in similar results (**Figure 2**). In this case 7 rabbits survived the treatment and the monitoring period, while one rabbit succumbed to the infection two days after treatment termination. However, in this case all the surviving rabbits developed protective immunity, surviving a SC challenge 14 days from infection. This protective immunity indicates that the spores germinated, secreted toxins and established an infection, enabling the host to initiate a specific immune response. This could be a result of the late treatment initiation, low antibiotic dose or both.

As for treating symptomatic animals, cefazolin was effective in treating rabbits with bacteremia of up to 10^4^ CFU/ml (experiment limitation). All attempts to use cefazolin for the treatment of meningitis failed including very high dose of 200 mg/ml and boosting the treatment with direct injection of the antibiotic into the CSF, bypassing the need for crossing the BBB. Altogether, according to the rabbit model, cefazolin can serve as an alternative for the first line antibiotics in case of resistance but only as a PEP or for early-stage patients.

The combination of meropenem and doxycycline, the CDC first choice treatment for systemic anthrax patients is highly effective, at least according to the rabbit pulmonary model [31, 36]. Since we have already established that doxycycline is effective as a monotherapy under these conditions [32], the most important finding is that these substances do not inhibit each other. According to our previous findings meropenem was effective in treating CNS infections only in high doses [30, 31]. However, the dosage in the CDC guidelines was derived from experience in treating meningitis caused by different bacteria [28], that would likely be sufficient for *B. anthracis*, especially since the treatment includes a tetracycline (minocycline or doxycycline) with proven efficacy [32, 33].

## Material and Methods

### Bacterial strains, media and growth conditions

*B. anthracis* Vollum strain (ATCC14578) [40] was used in this study. Spores were used for IN instillation or nasal sprays. For CNS infections, *B. anthracis* spores were germinated by incubation in Terrific broth (Merck T9197) for 30 min followed by 2h incubated in DMEM-10% NRS at 37^°^C in 10% CO_2_ atmosphere to induce capsule formation.

### Infection of rabbits

New Zealand white rabbits (2.5-3.5 kg) were obtained from Charles River (Canada). The animals received food and water ad libitum.

#### Intranasal instillation (PEP)

Rabbits were anesthetized using 100mg ketamine and 10mg xylazine and infected by pipetting small drops of spore suspension into their nasal cavities with a 1 ml tip. The infection dose was 1×10^6^ CFU of spores (10xLD_100_) in a total volume of 1 ml. Animals were treated with antibiotics 6 -24 hours post inoculation (as described in the results).

#### Respiratory infection

Rabbits were anesthetized using 100mg ketamine and 10mg xylazine and infected by spraying spore suspension into their nasal cavity with a MicroSprayer® Aerosolizer for Rat — Model IA-1B-R (PennCentury™). The infection dose was 1×10^4^ CFU of spores (10xLD_100_). Following infection, the instrument was visually inspected for signs of blood, which might imply nasal cavity injury and direct blood stream spore deposition. In all cases the device was confirmed negative. Blood samples were drawn from the rabbits’ ear veins to determine bacteremia by total viable counts 16 hours post inoculation (CFU.ml-1) and the animals were immediately treated with antibiotics (as described in the results).

#### CNS infection

For ICM administration, the animals were anesthetized using 100mg ketamine and 10mg xylazine. Using a 23 G blood collection set, 300 μl of encapsulated vegetative bacteria were injected into the cisterna magna. The remaining sample was plated for total viable counts (CFU.ml-1). The animals were observed daily for 14 days or for the indicated period.

### Antibiotic treatment

Following infection, animals were treated as described in the results. Antibiotic doses were as follows: Doxycycline 15 mg/kg (Doxycycline-ratiopharm SF 100 mg/5ml), Meropenem 40 or 150 mg/kg (Anfarm 500 mg I.V.), Cefazolin 30 or 60 mg/kg (Cefazolin Panpharma 1g). The MIC for the Vollum strain; Doxycycline is 0.016-0.032 μg/ml (Etest), Meropenem is 0.064 μg/ml (by Etest) and Cefazolin is 0.15 μg/ml (microdilution).

### Ethics

This study was carried out by trained personnel, in strict accordance with the recommendations of the Guide for the Care and Use of Laboratory Animals of the National Research Council. All protocols were approved by the IIBR committee on the Ethics of Animal Experiments. We used female rabbits in these experiments since there are no significant differences in *B. anthracis* pathogenicity between male and female rabbits [41]. Before inoculation, animals were sedated using 100mg ketamine and 10mg xylazine. Animals were monitored twice a day and euthanized immediately by a 120 mg/kg sodium pentobarbitone injection when one of the following symptoms was detected: severe respiratory distress or the loss of righting reflex. Animals unable or unwilling to drink were injected with 20-100ml of saline or dextrose isotonic solution SC. Since in most cases anthrax symptoms are visible only in close proximity to death, there were cases where animals succumbed to the disease.

### Statistical analysis

The significance of the differences in survival rates between treated groups and untreated controls and of the differences in bacteremia and time to death were determined by Log-rank, using Prism 6 software (GraphPad, USA).

